# Hierarchical coding of local and global positions in episodic memory for naturalistic sequences

**DOI:** 10.1101/2025.08.26.672267

**Authors:** Xiaojing Peng, Yu Zhou, Liang Zhang, Tong Li, Paul S. Muhle-Karbe, Gui Xue

**Affiliations:** State Key Laboratory of Cognitive Neuroscience and Learning & IDG/McGovern Institute for Brain Research, Beijing Normal University, Beijing 100875, China; School of Psychology & Centre for Human Brain Health, University of Birmingham, Birmingham, United Kingdom; Chinese Institute for Brain Research, Beijing, China

## Abstract

Life unfolds continuously across multiple timescales, yet how the brain encodes hierarchical event sequences remains poorly understood. Using magnetoencephalography, we investigated the neural coding of hierarchical positions in naturalistic sequence memory. Participants viewed video sequences with a nested structure and later recalled their serial order. Behavioral analyses revealed transposition errors for items occupying identical global or local position. Multivariate classification and subspace analyses revealed separable neural representations of global and local positions. Local positions were better decoded in high-dimensional space, whereas global positions exhibited greater pattern separability in low-dimensional space. Brain regions encoding global positions exhibited longer neural timescales and reduced sensitivity to low-level event transitions compared to regions encoding local positions. These findings support a parallel hierarchical organization of content and context, a novel framework that enables integration of item and order across hierarchical levels and provides principles for hierarchical representations in both human memory and artificial intelligence.

**Highlights:**

- Hierarchical position coding in sequence memory for naturalistic stimuli
- The neural representations of global and local positions are separable
- Global and local position codes are low and high dimensional, respectively
- Global position regions have longer timescales than local position regions

## INTRODUCTION

Imagine a typical scenario of visiting a large zoo. You start your day in the reptile house, move on to the aviary, and end in the big-cat enclosure. Later, when recounting the trip to a friend, you not only remember the animals you saw but also the correct temporal order: reptiles before birds before lions. This ability to organize experiences into the right sequence is fundamental to daily life and underpins cognitive functions such as motor control, language comprehension, spatial navigation, and episodic memory (Dehaene et al., 2015; Yokoi & Diedrichsen, 2019).

For simple linear sequences, converging evidence suggests that the brain does not store order by directly chaining adjacent items. Instead, it employs a factorized representation that separately encodes event content and temporal position (Behrens et al., 2018; Fan et al., 2024; Logan & Cox, 2023; Osth & Hurlstone, 2023). This scheme accounts for behavioral phenomena such as transposition errors between items in the same within-list (Conrad, 1960; Osth & Dennis, 2015) or within-group position (Hensen, 1999; Ng & Mayberry, 2002), the flexible reconfiguration of sequences (Liu et al., 2019; Vaswani et al., 2017), and the transfer of learned structure to novel content (Fan et al., 2021; Whittington et al., 2020). In addition to behavioral and modeling evidence supporting position coding (Brown et al., 2000; Hensen, 1998; Peng et al., 2025), positional information is found in the phase of theta oscillations (Bahramisharif et al., 2018; Lisman & Jensen, 2013), in distributed activation within the hippocampus (Hsieh et al., 2014; Liu et al., 2022), and in prefrontal cortical neural subspaces that enable flexible order manipulation in working memory (Chen et al., 2024; Xie et al., 2022).

Real-world experiences, however, often contain hierarchical temporal structures. A zoo visit can be divided into thematic “chapters” (reptiles, birds, mammals), each containing a sequence of finer-scale events. Behavioral and neural evidence indicates that the brain can encode both local positions (e.g., second animal in the reptile house) and global positions (e.g., first section of the trip) within the same episode (Brown et al., 2000; Fan et al., 2024; Henson & Burgess, 1997; Kikumoto & Mayr, 2018; Shpektor et al., 2024). For example, local and global positions have been shown to be represented through theta- and alpha-band neural spatial patterns (Kikumoto & Mayr, 2018), in orthogonal neural space (Fan et al., 2024), and along the posterior to anterior axis of entorinal cortex (Shpektor et al., 2024). Despite these advances, the representational principles that enable hierarchical coding, and the mechanisms that bind temporal positions to event content, remain poorly understood.

One fundamental question is whether local and global positions are encoded within shared or distinct neural substrates. Shpektor et al. (2024) reported a posterior-anterior gradient in the entorhinal cortex, with positional coding shifting from lower- to higher-level, potentially paralleling the progression of spatial receptive fields along the same axis. In contrast, Fan et al. (2024) observed local and global positions are represented by orthogonal dimensions within overlapping prefrontal and temporoparietal regions. Importantly, neither approach directly addresses how positional information is bound to event content across hierarchical levels. According to the hierarchical process-memory framework (Hasson et al., 2015), cortical regions integrate information over its preferred timescale before passing it to higher-order areas (Chang et al., 2022). This principle is mirrored in hierarchical event segmentation (Kurby & Zacks, 2008; Zacks et al., 2007) and the underlying neural substrates (Baldassano et al., 2017), suggesting that position-content binding may emerge from common anatomical and temporal constraints. Thus, a critical next step is to determine how the neural encoding of local and global positions within this hierarchy enables the efficient binding of content and context, a unifying principle essential for understanding hierarchical representations in the brain.

A second question concerns the structure of local and global position codes in neural representational space. High-level positions are expected to remain stable over extended time windows and to abstract away from transient details, making them less sensitive to noise and variability. Low-dimensional coding is well-suited for such abstraction, enhancing stability (Li & Curtis, 2023; Sapountzis et al., 2022; Wasmuht et al., 2018), supporting generalization (Aghajanyan et al., 2020; Bernardi et al., 2020; MacDowell & Buschman, 2020), and reducing neural costs (Eliasmith et al., 2012; Laughlin, 2001). In contrast, low-level positions may benefit from high-dimensional coding, which offers greater representational capacity, supports fine-grained discriminations, and allows for flexible recombination of details (Fusi et al., 2016; Rigotti et al., 2013; Tang et al., 2019). Testing whether such a dimensionality division exists for hierarchical positions could reveal how the brain balances stability and flexibility across representational levels.

A third question is whether these hierarchical position codes operate robustly in naturalistic continuous experience. Prior work with extensively trained sequences (Kikumoto & Mayr, 2018; Shpektor et al., 2024) risks encouraging associative chaining (Schapiro et al., 2012) rather than genuine, content-independent positional coding (Tsao et al., 2018). Likewise, sequences of syllables or words typically exhibit higher within-group than between-group transition probabilities (Fan et al., 2024), potentially biasing participants toward relying on statistical transitions (Dehaene et al., 2015; Ding et al., 2016; Henin et al., 2021) instead of abstract positional structures. Disentangling positional coding from transitional probability processing is thus essential for understanding how the brain constructs and maintains hierarchical temporal frameworks under ecologically valid conditions.

In the present study, we address these questions using a naturalistic video-watching paradigm in which short clips are grouped by thematic content into higher-level episodes. Behavioral analyses examine whether participants spontaneously reorganize events into a hierarchical structure, and magnetoencephalography (MEG) based classification and subspace analyses probe the neural representations of local and global positions. These approaches allow us to identify the cortical regions and representational formats that support hierarchical sequence representations, and to test how these codes integrate with event content to enable flexible sequence memory.

## RESULTS

### Enhanced serial position memory for structured video sequences

In the main experiment with MEG recording (Exp.1), 26 participants viewed nine video clips presented sequentially and were later asked to recall the serial position of pseudo-randomly selected clips (see Methods for details). Each clip sequence depicted three distinct themes (e.g., meerkats in a desert, penguins in snow, monkeys in trees), each with three thematically related clips shown consecutively (Figure 1A). As a control, 33 participants completed a similar task without MEG recording (Exp.2), in which the nine clips were presented in a randomized order. Participants were not informed of the underlying structure. We hypothesized that participants in Exp.1 would spontaneously group the clips, resulting in both better overall performance and distinct error patterns, compared to Exp.2.

**Figure 1.**
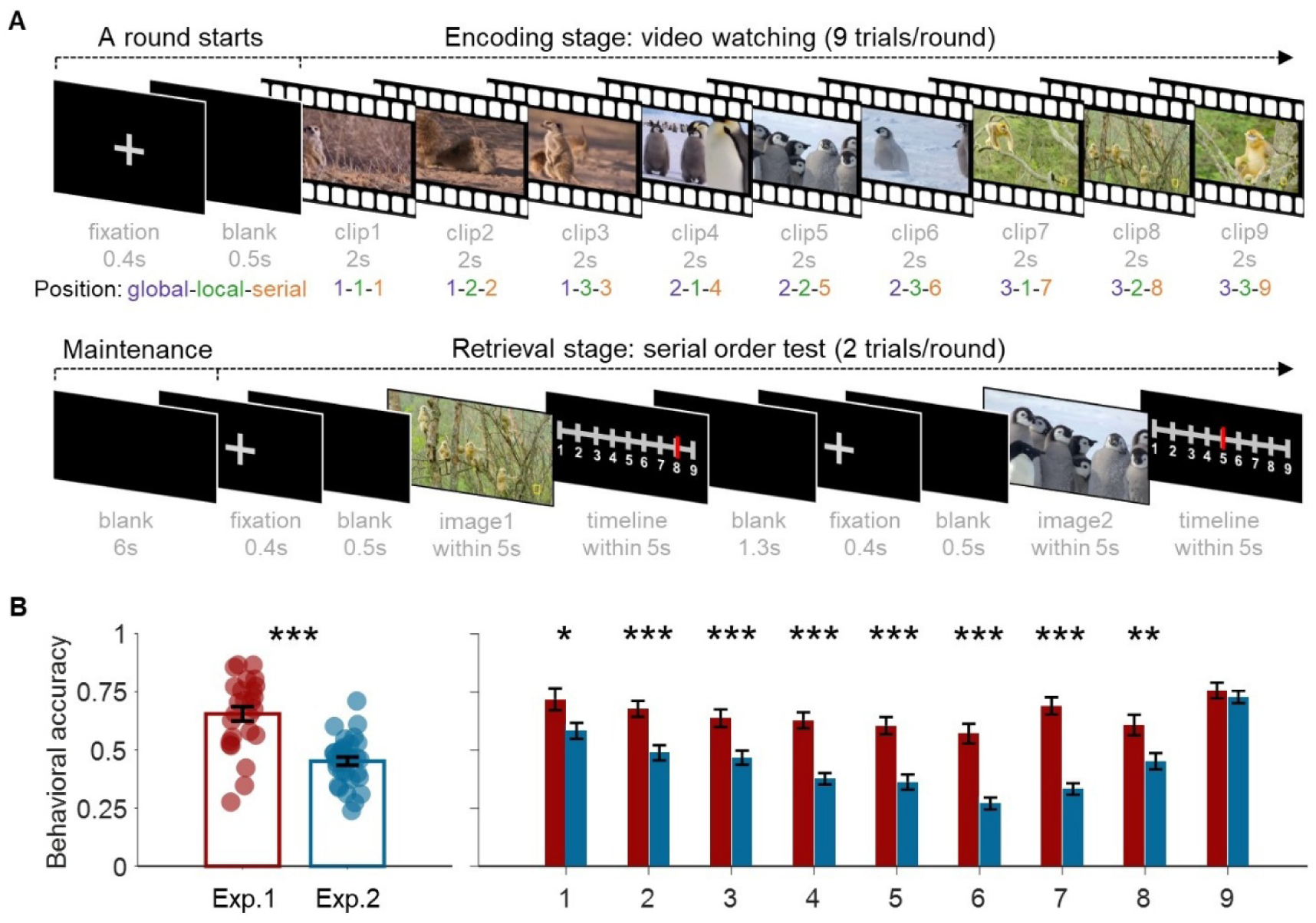
Experimental design and behavioral performance. (A) Task structure and stimuli organization. The schematic plot illustrates the learn-delay-test round used in Exp.1, where each round comprises nine video clips grouped by theme (i.e., animal). In contrast, these clips were presented in a randomized order in Exp.2. Both experiments included 90 unique sequences, with each serial position tested 20 times. (B) The overall accuracy (left) and position-wise accuracy (right) for both experiments. Error bars represent the standard error of the mean across participants. *p<0.05, **p<0.01, ***p<0.001.

Consistent with our hypotheses, accuracy was significantly higher in Exp.1 than in Exp.2 (65.47% vs. 45.13%; Figure 1B, left panel; 2×9 mixed-design ANOVA, main effect of experiment, *F*_(1,57)_ = 37.289, p < 0.001, 95% CI = [0.137, 0.270], partial *η*^2^ = 0.395). Simple effect analysis showed this advantage across serial positions 1 to 8 (Figure 1B, right panel; all p < 0.05; see Table S1 for details). These findings suggest that the hierarchical structure enhances serial order memory of the video sequences.

### Behavioral error patterns indicate hierarchical position coding

Drawing on prior research on serial order memory (Logan & Cox, 2023; Osth & Hurlstone, 2023; Polyn et al., 2009), we hypothesize that participants in Exp.1 would reorganize the sequence into a hierarchical structure, leading to more confusion errors among items sharing the same global or local position. In contrast, participants in Exp.2 are expected to show relatively more errors between adjacent serial positions.

To test these hypotheses, we constructed hypothetical confusion matrices reflecting different encoding strategies: global position coding, local position coding, and serial position coding (Figure 2A and 2B). In Exp.2, although the videos were presented in a randomized order, participants might still exhibit confusions between videos from the same theme. Accordingly, theme-based predictor was generated for each participant, based on their specific sequences. These predictors were then used to model individual error patterns. We applied generalized linear mixed modeling to quantify the unique variance explained (unique R^2^, see Methods) by each predictor.

**Figure 2.**
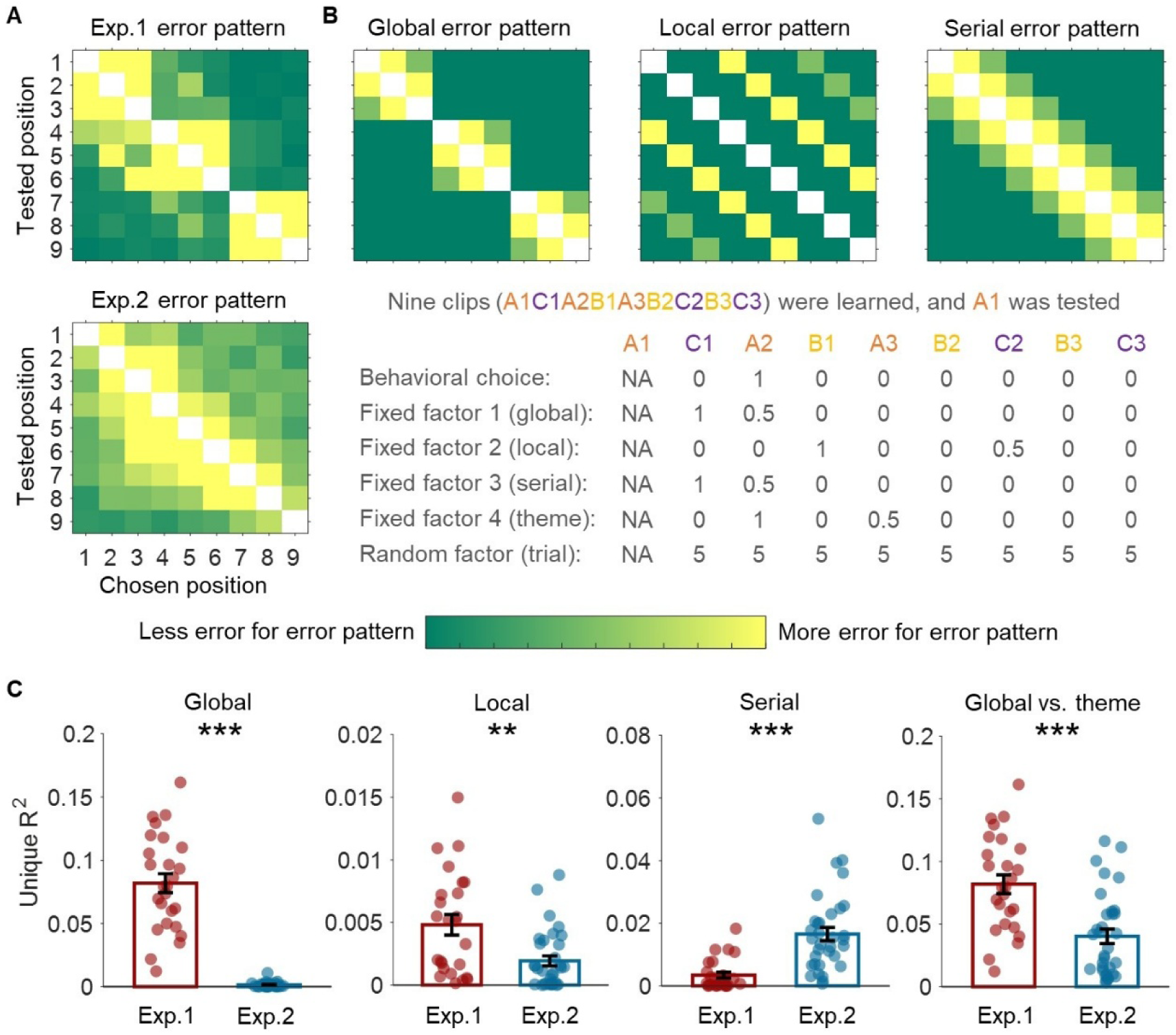
Behavioral error patterns indicate hierarchical position coding. (A) Group-averaged error patterns. (B) Hypothesized error patterns of different coding strategies. Top panel: The global, local, and serial error patterns are identical across Exp.1 and Exp.2. In Exp.2, a theme-based predictor is additionally included, individually defined for each trial and participant. Bottom panel: One example is shown. Letters A, B, and C represent distinct themes, and numbers 1, 2, and 3 indicate different video clips within the same theme. (C) Model fitting results. The analysis revealed distinct error profiles between the two experiments. Error bars indicate the standard error of the mean across participants. **p < 0.01, ***p < 0.001.

Supporting our hypotheses (Figure 2C), participants in Exp.1 showed significantly more confusion errors among items sharing the same global position (two-sided Wilcoxon rank-sum test: z = 6.54, p < 0.001, Cohen’s d = 3.18) and the same local position (z = 3.27, p = 0.0011, Cohen’s d = 0.62). In contrast, participants in Exp.2 exhibited more confusion errors between adjacent serial positions (z = -5.11, p < 0.001, Cohen’s d = 1.35). Furthermore, the unique R^2^ explained by global position coding in Exp.1 was significantly greater than that explained by theme-based coding in Exp.2 (z = 3.92, p < 0.001, Cohen’s d = 1.18), suggesting the combination of theme and temporal adjacency promotes chunking and the formation of hierarchical structure.

### Multivariate pattern analysis reveals hierarchical position coding

Having established behavioral evidence for global and local position coding in video sequences, we next examine whether local and global codes are reflected in distinct brain activity. At each encoding time point, we applied a one-vs-rest classification approach with five-fold cross-validation (see Methods), using whole-brain MEG spatial patterns as features. Cluster-based permutation tests (see Methods) revealed significant temporal clusters with above-chance decoding accuracy for both global and local positions (Figure 3A; all p < 0.05, cluster-corrected; see Figure S1 for decoding accuracy of each global and local position).

**Figure 3.**
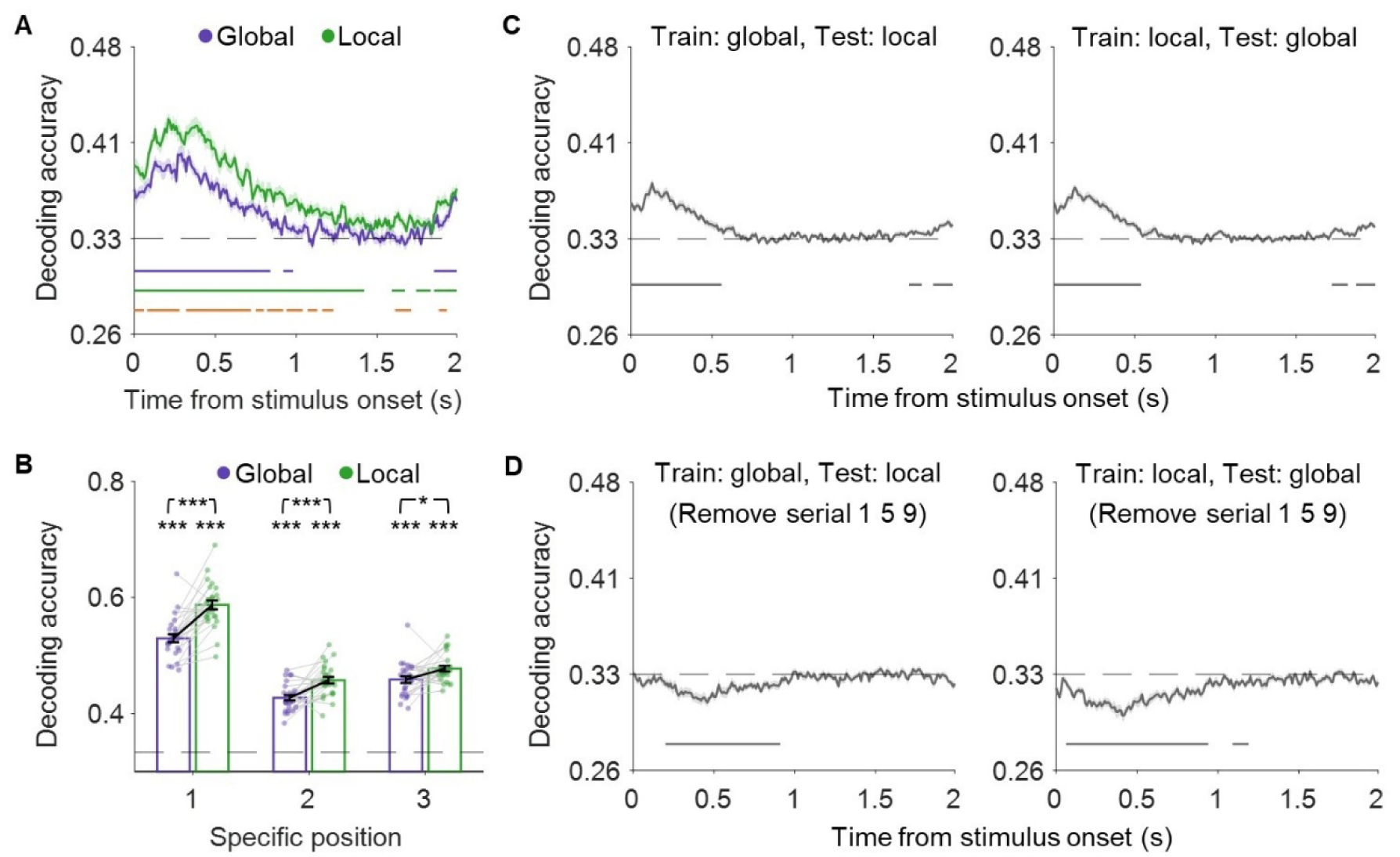
Multivariate decoding of local and global positions. (A) Decoding accuracy across time. Significant temporal clusters with above-chance decoding accuracy for global (purple) and local (green) positions are marked with colored lines. The gray dashed line indicates chance-level performance, and orange lines represent time points where decoding accuracy for local positions exceedes that for global positions. Shaded error bars represent the standard error of the mean across participants. (B) Decoding accuracy for each position. Error bars indicate the standard error of the mean across participants. *p < 0.05, ***p < 0.001. (C-D) Generalization between global and local positions (C) with all trials included and (D) with trials at serial positions 1, 5, and 9 removed from test dataset. Significant temporal clusters with above- and below-chance decoding accuracy between global and local positions are marked with gray lines. The gray dashed lines indicate chance-level performance. Shaded error bars represent the standard error of the mean across participants.

Repeated-measures ANOVA on the maximal decoding accuracy (Figure 3B, see Methods) revealed a significant effect of position (*F*_(2,50)_ = 233.89, p < 0.001, partial *η*^2^ = 0.903). Post hoc LSD tests further showed that position 1 was decoded more accurately than position 2 (t = 19.33, p < 0.001, 95% CI = [0.104, 0.128]) and position 3 (t = 15, p < 0.001, 95% CI = [0.078, 0.102]), consistent with the primacy effect (Peng et al., 2025; Pu et al., 2022; Xie et al., 2022). The effect of hierarchy was also significant, with higher decoding accuracy for local than global positions (*F*_(1,25)_ = 59.48, p < 0.001, partial *η*^2^ = 0.704, 95% CI = [0.026, 0.045]). We also found significant interaction between hierarchy and position (*F*_(2,50)_ = 7.47, p = 0.001, partial *η*^2^ = 0.23), suggesting the difference between local and global positions was higher in earlier position than in later positions. Overall, these results suggest that local positions are more distinctly represented in high-dimensional neural space.

To assess whether global and local positions are encoded by shared or distinct neural patterns, we trained classifiers on global positions to predict local positions, and vice versa (see Methods). This analysis revealed modest generalization between global and local positions (Figure 3C, all p < 0.05, cluster-corrected). However, when trials in which global and local positions matched (i.e., serial positions 1, 5, and 9) were excluded from test, no significant generalization was observed. If anything, performance fell below chance (Figure 3D; all p < 0.05, cluster-corrected). These results indicate that the representations of global and local positions are separable.

Finally, a generalized linear mixed model revealed that the neural representations of both global (t = 2.705, p = 0.0069, 95% CI = [0.33, 2.07]) and local (t = 2.427, p = 0.0153, 95% CI = [0.19, 1.81]) positions during encoding each uniquely predicted subsequent serial order memory performance (see Methods), suggesting the strength of global and local postion coding independently contributes to temporal memory.

### Local and global position coding in low-dimensional neural spaces

Previous work on working memory has shown that each local position is represented within an approximately orthogonal neural subspace, with increasing representational overlap for later positions (Xie et al., 2022). However, it remains unknown whether global position coding relies on similar neural mechanism, and whether global and local positions occupy shared or distinct neural subspaces.

To address these questions, we focused on the first 500 ms of the encoding period, during which peak decoding accuracy was observed for both global and local positions. We applied principal component analysis (PCA; see Methods) to reduce the dimensionality of whole-brain MEG data associated with each global and local position (Degutis et al., 2025; Li & Curtis, 2023; Murray et al., 2017). The first two principal components (PCs) accounted for 64%-73% of the variance within each position subspace and were used to define the neural subspaces for subsequent analyses.

To assess reliability of the estimated subspaces, we computed the ratio of variance explained (RVE; see Methods), which quantifies the variance captured when projecting data from one split onto the subspace derived from another split (Li & Curtis, 2023). Repeated-measures ANOVA revealed no significant main effect of hierarchy (*F*_(1,25)_ = 0.234, p = 0.633, partial *η*^2^ = 0.009) or hierarchy × position interaction (*F*_(2,50)_ = 2.441, p = 0.097, partial *η*^2^ = 0.089), indicating comparable reliability for global and local positions (Figure 4A). However, there was a significant main effect of position (*F*_(2,50)_ = 40.261, p < 0.001, partial *η*^2^ = 0.617). Post hoc LSD tests showed that position 1 exhibited greater representational stability than position 2 (t = 7.36, p < 0.001, 95% CI = [0.059, 0.103]) and position 3 (t = 6.46, p < 0.001, 95% CI = [0.056, 0.112]), suggesting greater representational reliability for early than late local and global positions.

**Figure 4.**
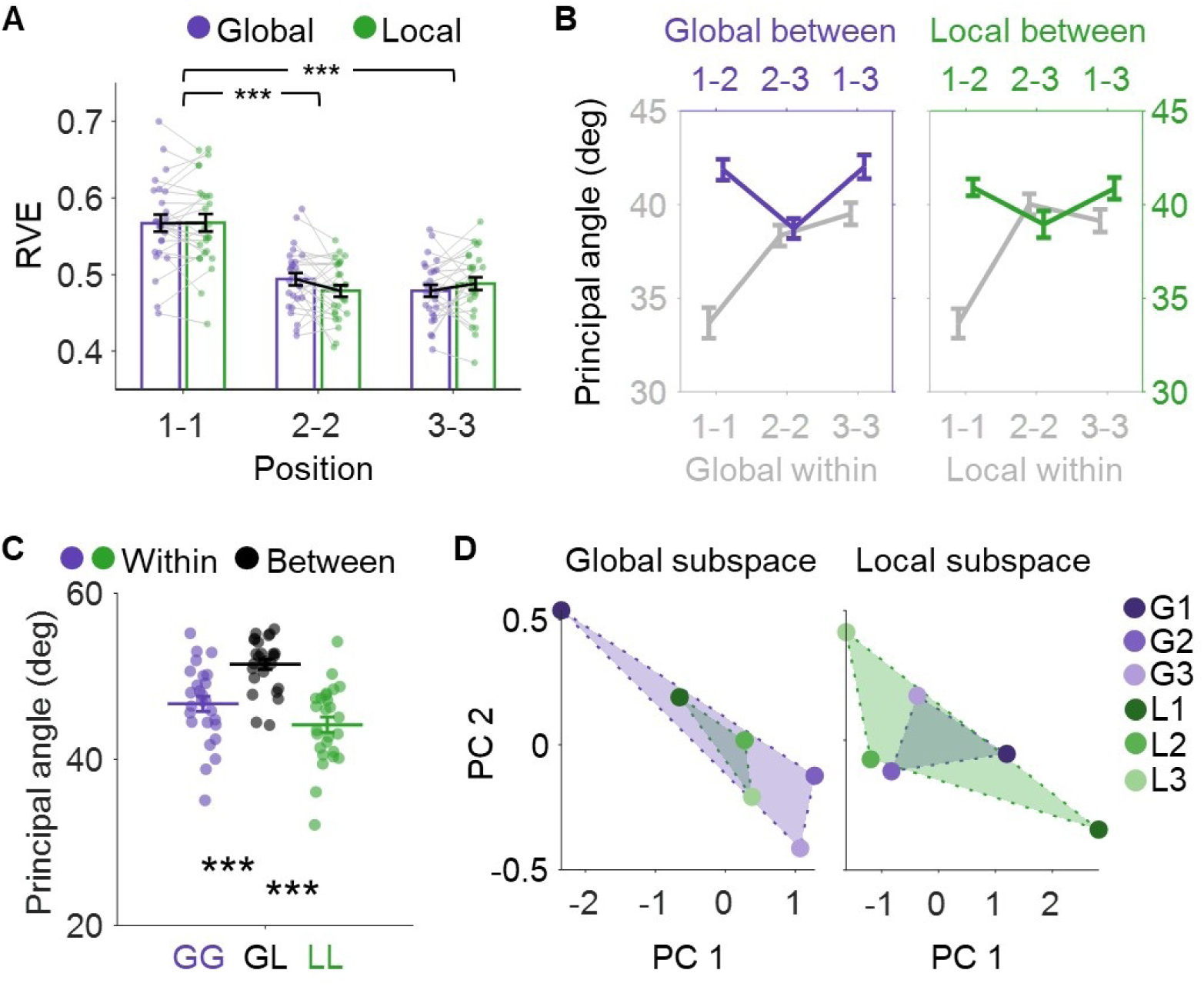
Neural subspace analysis for global and local positions. (A) Subspace stability measured by ratio of variance explained (RVE). Error bars represent the standard error of the mean across participants. ***p<0.001. (B) Subspace separability measured by principal angles. The top and bottom lines show between and within position principle angles, respectively. Error bars indicate the standard error of the mean across participants. (C) Principal angle between subspaces of global positions (GG, purple), between subspaces of global and local positions (GL, black), and between subspaces of local positions (LL, green). Error bars indicate the standard error of the mean across participants. ***p < 0.001. (D) Visualization of distinct local and global subspaces. The data were split into two halves. One subset was used to estimate global (or local) position subspace, while the data of global and local positions from the other subset were then projected onto the estimated subspace. G denotes global position and L denotes local position.

We further quantified the representational separability between positions using principal angles (see Methods) (Li & Curtis, 2023; Xie et al., 2022). The larger between-position relative to within-position angles indicate greater neural pattern separability. The analysis revealed significantly larger between-position than within-position angles, both between positions 1 and 2, and between positions 1 and 3, for both global and local positions (Figure 4B; all p < 0.03, permutation test; see Methods). In contrast, no significant separability was observed between positions 2 and 3, suggesting neural overlap for later positions. This decreasing neural differentiation complements the classification results (Figure 3B) and extends Xie et al. (2022)’s finding to indicate compressive coding for both global and local positions.

Crucially, our analysis revealed significantly greater subspace separability for global compared to local positions. Specifically, principal angles were larger for global than for local positions, both between positions 1 and 2 (p = 0.0125) and between positions 1 and 3 (p = 0.0095) (Figure 4B). Similar results were found using RVE (global 1-2 vs. local 1-2: p = 0.0093; global 1-3 vs. local 1-3: p = 0.0149; Figure S2A). These results contrast with the classification results in high-dimensional sensor space, where local positions showed higher decoding accuracy than global positions (Figure 3A and 3B). Together, these results suggest that global and local positions are better represented in low- and high-dimensional neural spaces, respectively.

To assess whether global and local positions are represented in shared or distinct neural subspaces, we computed the principal angle between their respective subspaces (see Methods, Figure 4C). This angle was significantly larger than that within global position subspaces (p < 0.001, permutation test) and within local position subspaces (p < 0.001). Similar results were observed for RVE (both p < 0.001, Figure S2B). For visualization, data from one split were projected onto subspaces estimated from another split (Figure 4D, see Methods). The variance of local positions 1, 2, and 3 shrank when projected onto global position subspace, and vice versa, indicating a shift in neural subspace (Li & Curtis, 2023). Together, these results corroborate the classification results to further suggest that local and global positions are encoded in distinct neural spaces.

### Distinct brain regions encode global and local positions

So far, both the classification and subspace analyses reveal distinct neural representations for global and local positions. To localize the sensors that contribute to the representations of hierarchical positions, we identified the top 25 sensors (including all sensors tied for 25th place) based on their frequency of significant contribution within the first 500 ms (see Methods) (Grootswagers et al., 2017; Haufe et al., 2014). The Dice similarity between global position (GP) and local position (LP) sensors was 0.296, indicating minimal overlap. GP and LP sensors were primarily located in the parietal region and temporal lobe, respectively (Figure 5A).

**Figure 5.**
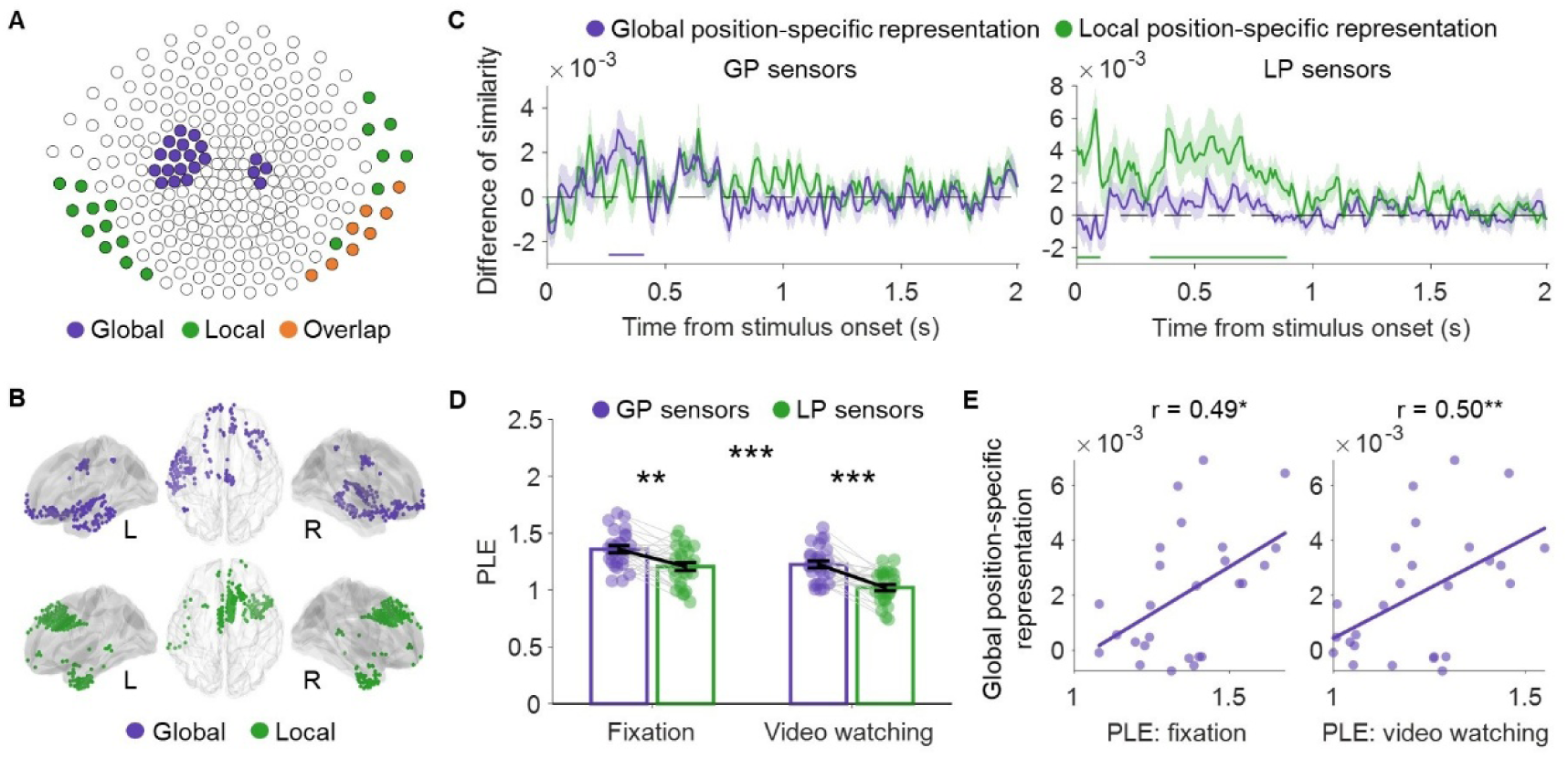
Brain regions encoding global and local positions show different neural timescales. (A) Global position (GP) and local position (LP) sensors. The top 25 sensors (including all sensors tied for 25th place) contributing to the decoding of global (purple) and local (green) positions were identified. Overlapping sensors (orange) were excluded for further analysis. (B) Source regions encoding global (purple) and local (green) positions. Overlapping vertices were excluded. (C) Dissociated position-specific representations of global (purple) and local (green) positions in GP and LP sensors. Neural pattern similarity was compared between trial pairs sharing the same global (or local) position and those with different positions. Temporal clusters showing significant difference of similarity are indicated by the respective colored lines. Shaded error bars represent the standard error of the mean across participants. (D) Power law exponent (PLE). PLE was calculated during fixation and video-watching periods for both GP and LP sensors. Error bars indicate the standard error of the mean across participants. **p < 0.01, ***p < 0.001. (E) Pearson correlation between PLE and global position-specific representation in GP sensors. *p<0.05, **p<0.01.

To identify brain regions involved in encoding global and local positions, we performed source reconstruction based on cortical meshes and projected the corrected classifier weights from sensor onto source space (see Methods) (Demarchi et al., 2019). GP vertices were primarily located in the rostral prefrontal cortex and temporal pole, whereas LP vertices were found in the posterior prefrontal cortex and inferior temporal regions (Figure 5B).

We further conducted representational similarity analysis (RSA) to verify that GP and LP sensors encode distinct positional information (Kriegeskorte et al., 2008) (Figure 5C; see Methods). A significant temporal cluster (270-420 ms) representing global positions was identified in GP sensors, whereas significant temporal clusters (0-110 ms and 320-900 ms) representing local positions were found in LP sensors (all p < 0.05, cluster-corrected). Additionally, significant temporal clusters representing serial positional information were found for both GP and LP sensors (Figure S3).

### GP sensors show longer neural timescales

Having identified dissociated brain regions representing global and local positions, we next examine how this functional dissociation aligns with hierarchical event representations. Specifically, we hypothesize that if the hierarchies for position and event content align, the regions encoding global positions would exhibit longer neural timescales and reduced sensitivity to low-level event transitions, compared to regions encoding local positions (Baldassano et al., 2017; Hasson et al., 2015; Wolff et al., 2022).

We computed the power law exponent (PLE; see Methods), which quantifies the temporal nesting of neural signals (i.e., how faster frequencies are embedded within more dominant slower ones), with larger PLE values indicating longer neural timescales (Wolff et al., 2019). As a sanity check, we first confirmed that the PLE during video-watching phase was significantly lower than that during fixation period (*F*_(1,25)_ = 191.23, p < 0.001, partial *η*^2^ = 0.88) (Figure 5D), consistent with previous findings (Golesorkhi et al., 2021). Critically, PLE was significantly higher for GP than LP sensors during both video watching (*F*_(1,25)_ = 21.83, p < 0.001, partial *η*^2^ = 0.47) and fixation (*F*_(1,25)_ = 12.65, p = 0.0015, partial *η*^2^ = 0.33), suggesting both intrinsic and task-related temporal properties were different between these regions. Further analysis revealed that GP sensors displayed more pronounced slow fluctuations, whereas LP sensors exhibited greater power in the high gamma range (Figure S4A). The PLE results were largely replicated when comparing the autocorrelation window (ACW) (Figure S4B), which measures the temporal decay of autocorrelation (Honey et al., 2012).

Finally, the PLE values of GP sensors during both fixation and video-watching periods were positively correlated with the strength of global position-specific representation (Figure 5E), further suggesting that longer neural timescales were associated with better global position coding.

### GP sensors show less sensitivity to low-level event transitions

To test whether GP and LP sensors are respectively sensitive to high- and low-level event content, we defined three boundary conditions (Figure 6A): (1) no boundary (within-event transitions), (2) low-level boundary (between events sharing the same theme), and (3) high-level boundary (between different themes). We then measured boundary-induced neural pattern shifts, quantified as multidimensional distance (MDD, see Methods) (Zheng et al., 2022).

**Figure 6.**
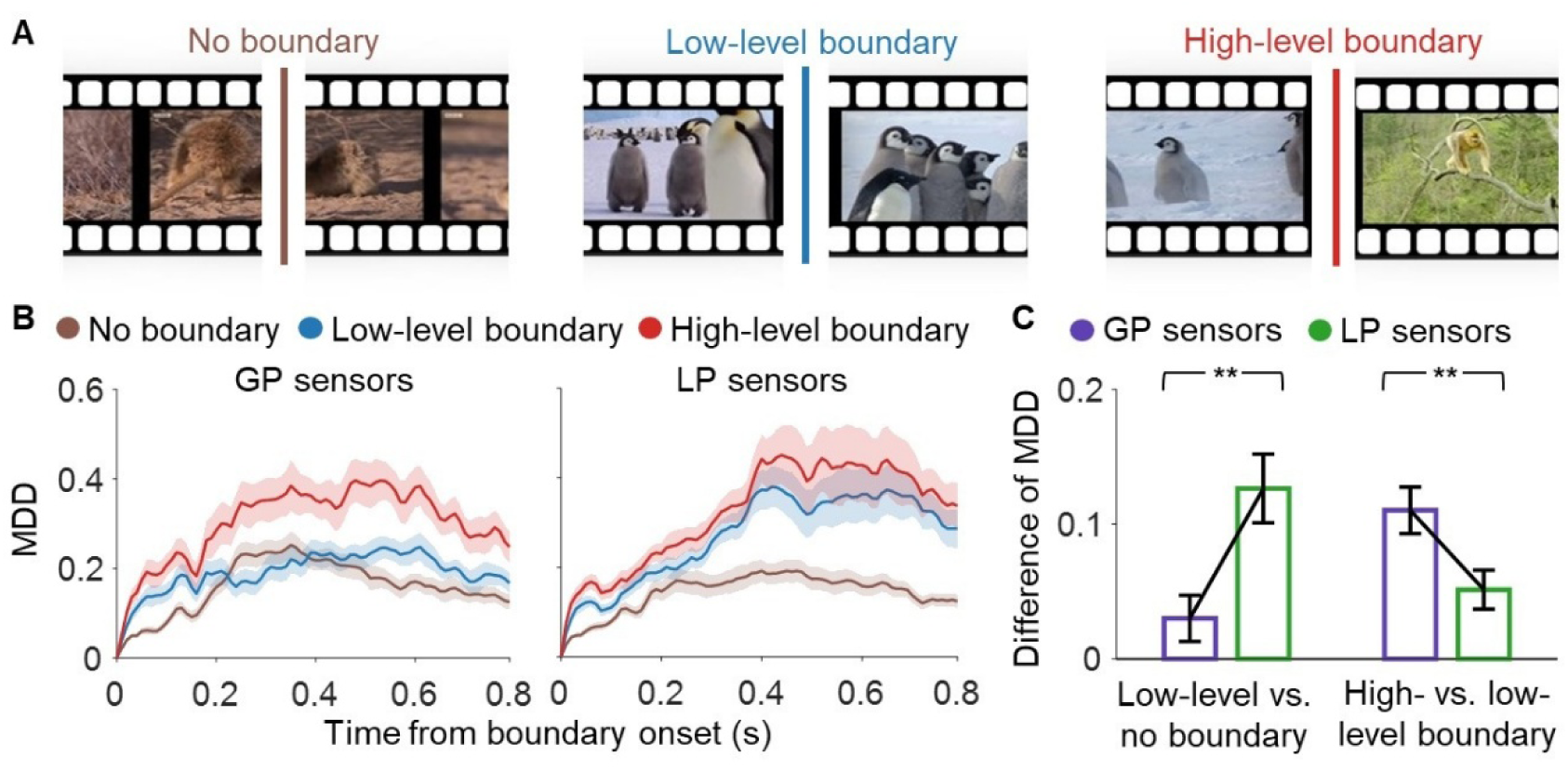
GP and LP sensors show sensitivity to different levels of event transitions. (A) Conditions of event boundaries. The no boundary (left), low-level boundary (middle), and high-level boundary (right) correspond to event transitions within the same video, within the same theme, and between different themes, respectively. (B) Multidimensional distance (MDD). Shaded error bars represent the standard error of the mean across participants. (C) Difference of MDD across boundary types and sensors. Error bars represent the standard error of the mean across participants. **p<0.01.

GP sensors exhibited pronounced MDD only after high-level boundaries, whereas LP sensors showed similar MDD after both low- and high-level boundaries (Figure 6B). We further contrasted low-level vs. no boundary and high-vs. low-level boundary to indicate the sensitivity to low- and high-level event transitions, respectively. Repeated-measures ANOVA revealed a significant hierarchy (GP vs. LP sensors) by boundary (high-vs. low-level) interaction effect (*F*_(1,25)_ = 24.85, p < 0.001, partial *η*^2^ = 0.50) (Figure 6C). Simple effect analyses showed that LP sensors were more sensitive to low-level event transitions (*F*_(1,25)_ = 13.64, p = 0.0011, partial *η*^2^ = 0.35), whereas GP sensors were more sensitive to high-level event transitions (*F*_(1,25)_ = 8.64, p = 0.007, partial *η*^2^ = 0.28).

## DISCUSSION

Our episodic experiences unfold as continuous streams but are perceived and remembered as hierarchically organized events. How the brain processes hierarchical temporal positions remains poorly understood. With a paradigm closely approximating real-life event structures, we provided converging behavioral and neural evidence that hierarchical position coding emerges spontaneously and relies on distinct neural representations. Global and local positions are better represented in low- and high-dimensional neural spaces, respectively, and are supported by dissociable brain regions. These differences mirror the anatomical hierarchy of event processing, as reflected in brain regions’ intrinsic neural timescales and sensitivity to event content.

Using a naturalistic paradigm, we observed robust evidence of hierarchical positional coding, consistent with prior behavioral and modeling results (Brown et al., 2000; Fan et al., 2024; Huttenlocher et al., 1990; Logan & Cox, 2023; Osth & Hurlstone, 2023). Specifically, participants made transposition errors when items shared identical global or local position, indicating reliance on multilevel positional structures to organize event sequences. Several features of the current study should be noted. First, it approximates real-world episodic memory with naturalistic stimuli. Second, by one-shot learning, it minimizes the across-item associative chaining and the influence of transition probability. Third, it includes a control condition, allowing the direct comparison of structured versus unstructured sequences. These methodological refinements collectively offer a more ecologically valid and rigorously controlled test of hierarchical position coding.

We found converging evidence for distinct neural representations of global and local positions using complementary approaches. Whole-brain classification revealed low generalization between global and local positions, and subspace analysis showed significant principal angle between them, indicating separable population codes. These results replicate and extend existing findings (Fan et al., 2024; Shpektor et al., 2024) to one-shot, episodic-like sequence memory. Unlike Fan et al. (2024), we found that the neural signals encoding global and local positions were localized to distinct brain regions, rather than orthogonalized representations within the left prefrontal and temporoparietal regions. This suggests that position coding during short-term working memory and long-term episodic memory might employ distinct coding schemes. Supporting this view, Shpektor et al. (2024) trained participants to memorize 113-tone auditory sequences through repeated listening and testing over 2 days, and found distinct voxel

populations within the entorhinal cortex encoding low- vs. high-level positions.

More importantly, we observed that brain regions encoding global positions exhibit longer neural timescales and reduced sensitivity to low-level event transitions relative to regions encoding local positions. These findings extend the hierarchical process-memory theory (Hasson et al., 2015) to incorporate both event content and position, and further suggest a unified organizational principle of the cortical hierarchy for processing different types of information. Within this unified neural hierarchy, the factorized coding scheme (Behrens et al., 2018) can be readily implemented at different hierarchical levels. A similar hierarchy has been observed in the entorhinal cortex, where an anterior-posterior anatomical gradient not only corresponds to a spatial gradient of grid cells but also encodes hierarchical temporal positions (Shpektor et al., 2024). Future studies using complementary imaging technologies and diverse paradigms can examine the hierarchical representations of temporal information in cortical regions (Liu & Buonomano, 2025; Paton & Buonomano, 2018) and the hippocampal-entorhinal circuit (Buzsáki & Moser, 2013; de Sousa et al., 2021; Eichenbaum, 2014; Tsao et al., 2022).

We provide the first empirical demonstration of a clear dissociation between local and global position representations, revealing their distinct dependence on high- and low-dimensional neural coding schemes, respectively. This distinction supports a fundamental principle of hierarchical factorization: higher-level, abstract representations adopt low-dimensional codes to promote stability and robustness. Specifically, global positions, which require integration over extended time windows, are encoded in low-dimensional codes that compress complex inputs into compact, generalizable representations (Aghajanyan et al., 2020; Bates & Jacobs, 2020; Bernardi et al., 2020; MacDowell & Buschman, 2020; Wasmuht et al., 2018). In contrast, local positions depend on high-dimensional codes to preserve fine-grained temporal details, facilitating discrimination, rapid learning, and episodic memory (Fusi et al., 2016; Rigotti et al., 2013; Sheng et al., 2022; Tang et al., 2019). Future studies should examine this principle in other domains, such as spatial context and event content.

Notably, we observed compressive coding for both local and global positions, as revealed by classification and subspace analyses. In particular, decoding accuracy and principal angles between position subspaces both decreased for later positions. These findings extend the results of Xie et al. (2022) to hierarchical position coding. This shared compressive coding pattern aligns with behavioral signatures in serial order memory, such as the primacy effect at different hierarchical levels (Fan et al., 2024). The compressive feature likely reflects an adaptive trade-off between representational efficiency and fidelity, allowing the brain to prioritize early positions within each hierarchy (Heusser et al., 2018; Pu et al., 2022), while also conserving resources for extended sequences.

These findings motivate a new framework: the parallel hierarchical organization of content and context (PHOCC) framework. This framework posits that the brain encodes event content (e.g., sub-events within episodes) and context (e.g., positions within temporal frameworks) in a layered, mutually aligned manner. For instance, the event “watching lizards” is nested within the broader content of “visiting a zoo”, which itself is part of “having a vacation”.

Simultaneously, its contextual setting, the “morning” is nested within the larger contexts of “Saturday” and “August”. This parallel, multi-scale representational system allows nodes in the content hierarchy to map directly onto corresponding levels in the context hierarchy, facilitating alignment across abstract levels (Baldassano et al., 2017; Kurby & Zacks, 2008; Zacks et al., 2007). This parallel structure is distinct from approaches that encode event content hierarchically but treat context in a linear and non-hierarchical manner (Buzsáki & Tingley, 2018; Horner et al., 2016; Howard & Kahana, 2002; Polyn et al., 2009; Pu et al., 2022).

The PHOCC framework offers multiple advantages. First, it enables multi-scale binding, allowing event content to be linked with their contexts at fine- and coarse-grained levels, thereby increasing retrieval flexibility (Polyn et al., 2009; Reagh & Ranganath, 2023; Wen & Egner, 2022). Second, it facilitates contextual inference, whereby missing details about either content or context can be reconstructed by navigating their corresponding hierarchies, consistent with Bayesian accounts of cognitive inference (Griffiths et al., 2008). Third, it supports schema formation across scales, enabling abstraction of recurrent structures at different levels (Baldassano et al., 2018; Reagh & Ranganath, 2023), such as a “typical zoo visit” and a “typical vacation”, and linking them together (Ghosh & Gilboa, 2014; Preston & Eichenbaum, 2013). Finally, by aligning segmentation boundaries in context and event content, this system reduces interference between similar memories and provides distinct retrieval cues, yielding more efficient encoding, organization, and transfer of knowledge (Howard & Kahana, 2002; Zacks et al., 2007).

These principles have major implications for artificial intelligence. Current systems often rely on flat representations, limiting their ability to generalize across contexts, leading to catastrophic forgetting (Kirkpatrick et al., 2017), and constraining compositional generalization and long-term reasoning (Lake et al., 2017). Parallel hierarchical organization offers a solution: by nesting event content and contexts across temporal and spatial scales, agents can disambiguate overlapping experiences, reconstruct missing information, and perform multi-scale inferences, analogous to contextual reinstatement in human memory (Folkerts et al., 2018; Manning et al., 2011). This approach underlies advances in hierarchical reinforcement learning (Botvinick et al., 2019; Vezhnevets et al., 2017) and generative world models (Franklin et al., 2020), providing the structural priors needed for compositional transfer and robust abstraction. By embedding parallel content-context hierarchies, artificial intelligence systems could achieve more adaptive, memory-efficient, and human-aligned learning.

In summary, by integrating a one-shot episodic sequence paradigm with naturalistic stimuli and novel analytical approaches, the present study reveal distinct neural coding of global and local positions that aligns with hierarchical event content coding. These findings motivate a parallel hierarchical organization of content and context (PHOCC) framework, which not only elucidates the neural mechanisms underlying human sequence memory but also provides a principled blueprint for designing artificial intelligence systems capable of hierarchical representation, multi-scale inference, and flexible generalization.

## DATA AVAILABILITY

The data and code that support the findings of the study are available from the corresponding author upon reasonable request.

## ACKNOWLEDGEMENTS

The authors thank Lei Wang and Sze Chai Kwok for the discussion at the early stage of this project, and Nikolai Axmacher for helpful comments on the manuscript. This work was supported by the National Natural Science Foundation of China (32441110 and 32330039).

## AUTHOR CONTRIBUTIONS

Conceptualization: X.P., P.S.M.-K., and G.X. Methodology: X.P., Y.Z., L.Z., T. L., P.S.M.-K., and G.X. Investigation: X.P. Formal analysis: X.P. Visualization: X.P. Funding acquisition: G.X. Writing – original draft: X.P. and G.X. Writing – review & editing: X.P., P.S.M.-K., and G.X.

## METHOD DETAILS

### Participants

In Exp.1, 26 volunteers (7 male) aged between 19 and 28 years (mean age: 21.69 years) participated. In Exp.2, 33 participants (7 male) aged between 19 and 26 years (mean age: 21.88 years) were recruited. All participants were right-handed, had normal or corrected-to-normal vision, normal color perception, and no history of neurological or psychiatric disorders, and were not on hormonal medications prior to testing. Participants were native Chinese speakers and recruited from Beijing Normal University and surrounding areas. The study was approved by the Ethics Council of Beijing Normal University (ICBIRA0072013), and all participants provided written informed consent and received financial compensation for their participation. No prior studies have directly compared the neural representations of global and local positions, so we could not predetermine sample sizes. However, our sample sizes are consistent with those in related studies (Michelmann et al., 2016; Michelmann et al., 2019).

### Materials

The stimulus materials were obtained from the Internet and manually edited using Adobe Premiere Pro CS6, then exported in a uniform output format via Adobe Media Encoder CS6. Each video clip was 2 s long, devoid of camera cuts, subtitles, or audio, with a resolution of 1280 × 720 pixels and a frame rate of 30 fps. The video clips primarily featured activities from 15 animal species across three distinct scenes (i.e., snowy field: bird, polar bear, wolf, penguin, and red fox; green forest: kangaroo, horse, squirrel, giraffe, and monkey; yellow plain: lion, ibex, meerkat, elephant, and zebra). In total, 810 unique video clips were used, with 54 clips for each theme.

### Experimental design and procedure

Each round consisted of three phases: encoding, maintenance, and retrieval (Figure 1A). During the encoding phase, after a 0.4-s fixation cross and a 0.5-s blank screen, participants were sequentially presented with nine unique clips from three themes (three clips per theme) and instructed to remember their serial order. Following a 6-s maintenance period with a blank screen, participants performed two retrieval tests. In each test, a static frame extracted from a randomly chosen clip was shown, and participants were asked to recall the serial position of the corresponding clip. They pressed a button with their left index finger as soon as they had an answer. A timeline was then displayed, and participants used their right index and middle fingers to move a red cursor to the correct position within 5 s, and confirmed their response by pressing the button with left index finger. The initial position of the cursor was randomized for each test.

For each round, the nine video clips were from three themes, with the three clips per theme differing in fine-grained details (e.g., specific animals, scenes, and activities). In Exp.1, the nine clips followed a nested structure (i.e., clips from the same theme were presented consecutively), whereas Exp.2 employed a fully randomized order. The two test frames were always drawn from two different themes. To minimize interference, no video clip was repeated across the entire experiment, and the same theme did not reappear within at least four subsequent rounds. The experiment consisted of 90 trials, evenly divided into 10 runs. Each serial position was probed 20 times in total.

### Data collection

The MEG experiment (Exp.1) was programmed using MATLAB R2014b (The MathWorks, Inc.) and Psychtoolbox 3.0.18, running on a 64-bit Windows 7 platform. Neurophysiological data were collected in a magnetically shielded room using a 275-channel CTF MEG system at the Institute of Biophysics, Chinese Academy of Sciences, Beijing, China. Due to sensor malfunctions (LF55, RT16, and RT23), data from 272 sensors were recorded. Signals were sampled at 1200 Hz throughout the experiment. To localize participants’ head positions relative to the MEG sensors, three localization coils were placed at the left and right preauricular points and the nasion. To improve source localization, individual T1-weighted brain images were acquired using a Siemens 3T Prisma scanner at the Beijing Normal University MRI Center, Beijing, China. Exp.2 was conducted outside the MEG scanner at Beijing Normal University.

### Analysis of behavioral error patterns

The true error patterns were assessed based on the averaged swap errors between positions. For hypothesized confusion matrices, we used values of 1, 0.5, and 0 to denote high, intermediate, and low probabilities of swap errors.

For example, in global position coding, position 1 would have a high probability of swapping with position 2, an intermediate probability of swapping with position 3, but would be unlikely to swap with positions 4 to 9. Similarly, for local position coding, position 1 would have a high probability of swapping with position 4, an intermediate probability of swapping with position 7, but would be unlikely to swap with positions 2, 3, 5, 6, 8, or 9. For serial position coding, position 5 would have a high probability of swapping with positions 4 and 6, an intermediate probability of swapping with positions 3 and 7, but would be unlikely to swap with positions 1, 2, 8, or 9.

Additionally, participants in Exp.2 might confuse videos from the same theme. The theme-based predictor was constructed individually for each participant and each trial, with values of 1 and 0.5 indicating that a position belonged to the same theme at a closer or more distant serial position, respectively, and 0 indicating that it belonged to a different theme.

Following a previous study (Kietzmann et al., 2019), the empirical error patterns in Exp. 1 were modeled using three candidate error patterns (global, local, and serial) within a full generalized linear mixed model (GLMM) that incorporated a random intercept to account for trial-level variance. The unique variance explained (unique R^2^) for each predictor was calculated by subtracting the variance explained by a reduced GLMM (excluding the predictor of interest) from that of the full GLMM. This procedure was performed separately for each predictor and each subject. In Exp. 2, one additional predictor (i.e., the theme-based predictor) was estimated.

### Preprocessing of MEG data

All analyses were conducted in MATLAB R2020b (The MathWorks, Inc.) using the FieldTrip toolbox (Oostenveld et al., 2011) and custom scripts.

Preprocessing followed standard procedures outlined in a previous study (Ferrante et al., 2022). To remove power-line noise, sensor signals were filtered using a fourth-order Butterworth IIR filter, targeting a stopband of 49.5-50.5 Hz and its harmonics (99.5-100.5 Hz, 149.5-150.5 Hz, 199.5-200.5 Hz, and 249.5-250.5 Hz). To remove slow drifts, a high-pass filter at 0.5 Hz (fourth-order Butterworth IIR) was applied. Eye-movement and heartbeat artifacts were eliminated via independent component analysis. Epochs containing jump or muscle artifacts were detected using FieldTrip’s auto-algorithms and excluded from further analysis.

For classification and subspace analyses, the signal-to-noise ratio (SNR) was improved by smoothing the preprocessed MEG data with a 25-ms sliding window. The data were demeaned across sensors at each time point per participant to remove global signal fluctuations (e.g., sensory adaptation or system-wide artifacts) that uniformly affect all sensors (Kalm & Norris, 2017; Kikumoto & Mayr, 2018). In addition, mean activity across trials was subtracted for each sensor and time point to eliminate trial-invariant patterns (e.g., stimulus-driven baseline shifts) (Fan et al., 2021; Grootswagers et al., 2017). Finally, the data were downsampled to 100 Hz to improve computational efficiency (Grootswagers et al., 2017).

### Classification analysis

Lasso-regularized logistic regression models were employed to decode global and local positions (Liu et al., 2019), using one-vs-rest classification and five-fold cross-validation. For each participant, position, and encoding time point, binary classifiers were trained using preprocessed whole-brain sensor data. The predicted label for each trial/time point was determined by the classifier yielding the highest evidence among all three classifiers (Liu et al., 2019). Correct classifications were coded as 1 and incorrect classifications as 0. Decoding accuracy at each time point was quantified as the proportion of trials with correct classification (Figure 3A, 3C, and 3D; Figure S1).

To compare decoding accuracy between global and local positions (Figure 3B), we extracted the maximal decoding accuracy for each participant and condition within their significant temporal clusters (Figure S1).

For cross-classification between global and local positions (Figure 3C, 3D), we performed split-half cross-validation. In an additional analysis (Figure 3D), serial positions 1, 5, and 9 were excluded from the test dataset because they share the same local and global positions. These procedures were repeated 30 times, and decoding accuracies from the 30 iterations were averaged to obtain reliable results.

### Subsequent memory effect

To test whether the neural representations of both global and local positions during encoding could predict subsequent serial order memory, decoding accuracy for each trial was defined as the proportion of correctly classified time points across the encoding period. GLMM was then conducted to predict trial-level memory performance (i.e., correct or incorrect) based on the strength of global and local position coding, as specified by the following formula:

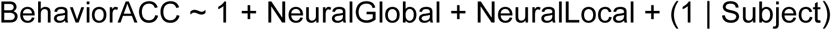

where the term (1 | Subject) specifies a random intercept for subjects. The GLMM was implemented using the MATLAB function fitglme.

### Localization of brain regions representing local and global positions

#### Weight correction

As classifiers can assign nonzero weights to sensors that contain no class-specific information, uncorrected (raw) weights cannot be interpreted directly (Grootswagers et al., 2017; Haufe et al., 2014). To address this issue, slope coefficients of the classifiers were corrected using the procedure from Haufe et al. 2014. Corrected weights (denoted as **W**_corrected_) associated with each global and local position were obtained by multiplying the feature covariance with the raw weights of the classifier:

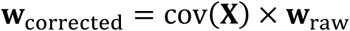

where **X** represents an N×M matrix of MEG power data, with N denoting the number of trials and M the number of features (i.e., 272 sensors), and **W**_raw_ denotes a length-M weight vector comprising the slope coefficients of the trained model. Each iteration of training and testing produced a set of slope coefficients associated with each classifier.

For each participant and each encoding time point, corrected weights, obtained from iterations of five-fold cross-validation, were averaged separately for each representation (e.g., global positions 1, 2, and 3). Sensors consistently contributing to the classification across all positions and participants were identified, defined as those whose corrected weights systematically deviated from zero according to Student’s t-tests and survived FDR correction. For each sensor, we computed its frequency of significant contribution within the first 500 ms of the encoding period, during which peak decoding accuracy was observed for both global and local positions. The top 25 sensors (including all sensors tied for 25th place) were identified as GP sensors (20 sensors after excluding 8 overlapping sensors) or LP sensors (18 sensors after excluding the same 8 overlapping sensors).

#### Source reconstruction

For the construction of individual head models, the anatomical MRI of each participant was first manually co-registered to the CTF coordinate system based on three fiducial landmarks: the nasion and the left and right preauricular points. Subsequently, the anatomical images were segmented into different tissue types, separating gray matter, white matter, and cerebrospinal fluid. Using the resulting brain mask, the single-shell method was then conducted to build individual head models.

To obtain individual source models, we employed FreeSurfer to reconstruct the cortical sheet. This procedure yielded a triangulated cortical mesh, ideally consisting of a number of approximately equally sized triangles that form a topological sphere for each of the cerebral hemispheres. Since the original FreeSurfer meshes typically contain more than 100,000 vertices per hemisphere, which is an excessive number for MEG source analysis, we downsampled them to 8,004 vertices with Connectome Workbench. This ensured both a faithful surface topology and low variance in triangle size. Finally, individual source models were aligned with the CTF coordinate system using FieldTrip.

Once the head and source models were obtained, we used FieldTrip to compute the individual leadfield matrix, a procedure known as forward modeling. Based on this, we estimated the unknown sources underlying the measured MEG signals, a step referred to as inverse modeling.

To identify GP and LP vertices, we first computed linearly constrained minimum variance (LCMV) spatial filters for each representation (e.g., global positions 1, 2, and 3) from the corresponding preprocessed sensor data, using the previously derived leadfield matrix. The sensor-level corrected weights obtained from the classification analysis, which were averaged across five-fold training and testing iterations, were then multiplied with the LCMV filters to generate informative activity patterns for each representation and each encoding time point (Demarchi et al., 2019).

Next, following the same procedure used to identify GP and LP sensors, we determined vertices that consistently deviated from zero across participants. Contributing vertices were defined as those surviving FDR-corrected Student’s t-tests across all representations. We restricted our analysis to the first 500 ms per vertex and quantified the frequency of significant contributions. Finally, the top 600 vertices (ties included) were selected, yielding 170 GP vertices (607 – 437) and 183 LP vertices (620 – 437) after excluding the 437 overlapping vertices.

### Subspace analysis

PCA (principal component analysis) was employed to identify low-dimensional subspaces encoding the conditions of interest. For each participant and each encoding time point, eigen decomposition was performed on the matrix **S**, structured as C (conditions) by M (sensors), representing whole-brain sensor activity patterns (i.e., preprocessed signals) averaged across trials of the same condition. **S** had column-wise zero mean, as the mean of each column was removed. Eign decomposition was conducted by the following formula (Degutis et al., 2025; Li & Curtis, 2023; Murray et al., 2017; Sapountzis et al., 2022):

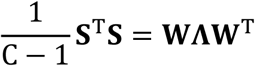

where each column of **W** was a unit-length eigenvector containing the weights of sensors on the principal component (PC) and the diagonal elements of **Λ** were the corresponding eigenvalues.

When estimating subspaces for each global and local position (Figure 4A and 4B, Figure S2A), nine conditions were included for each subspace. For the global position subspaces, trials corresponding to each global position were grouped into nine conditions, corresponding to all combinations of three local positions and three scenes (i.e., local position 1-3 × scene 1-3). This approach was adopted because the experimental design ensured that all three scenes appeared in each round, but could not guarantee that all 15 animals were evenly distributed across global positions 1, 2, and 3.

Consequently, grouping trials by scene yielded more balanced trial counts for each global position, helping to maintain a comparable SNR ratio. Specifically, trials from each scene were split into three groups corresponding to local positions 1, 2, and 3, and then averaged. This procedure can be viewed as providing multiple data points per sensor for the computation of the sensor-by-sensor covariance matrix (i.e., 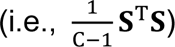, thereby improving the stability of the covariance estimation for each global position subspace. Similarly, for each local position subspace, the nine conditions included global positions 1 to 3 combined with scenes 1 to 3 (i.e., global 1 - scene 1, …, global 3 - scene 3).

To allow for more direct comparison between global and local position subspaces (Figures 4C,D and S2B, especially the visualization in Figure 4D), we defined the subspaces using the conditions corresponding to positions 1, 2, and 3. Specifically, the conditions of global position subspace estimation included global positions 1 (i.e., serial positions 1, 2, and 3), 2 (i.e., serial positions 4, 5, and 6), and 3 (i.e., serial positions 7, 8, and 9). Similarly, the conditions of local position subspace included local positions 1 (i.e., serial positions 1, 4, and 7) to 3 (i.e., serial positions 3, 6, and 9).

Throughout the study, we defined subspaces using the first two PCs with the largest eigenvalues, based on the weight matrix **T** (derived from the weight matrix **W**), which was of size M (sensors) by 2 (PCs).

### Visualization of subspaces

For visualization purposes (Figure 4D), sensor data from all subjects were concatenated, and PCA was applied to the aggregated sensor space (Degutis et al., 2025; Li & Curtis, 2023), with **S** constructed on the averaged encoding data from the first 500 ms. For example, the MEG data matrix **S**_global_, with a size of 3 (global positions 1, 2, and 3) by 7072 (272 sensors × 26 participants), was multiplied with the weight matrix T_local_, which contained the two columns with the largest eigenvalues in W_local_. The result of **S**_global_ × **T**_local_ was the projection of sensor activity pattern of each global position onto the local position subspace, commonly referred to as PC scores within the framework of PCA (Li & Curtis, 2023). The data were divided into two halves: one half was used to estimate the global and local position subspaces, while the data of global and local positions from the other half were then projected onto the estimated subspaces.

### Principal angle

To quantify the alignment between two separate estimates of subspaces, principal angle (Li & Curtis, 2023) was computed by applying singular value decomposition to the matrix 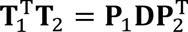, where **T**_1_ and **T**_2_ are weight matrices corresponding to the two separate subspaces, both sized 272 (sensors) by 2 (PCs). The matrix **D** was a diagonal matrix, with diagonal elements representing the ranked cosines of principal angles θ_1_ and θ_2_: D = diag(cos(θ_1_), cos(θ_2_)). The first principal angle was reported and statistically tested (Li & Curtis, 2023; Xie et al., 2022). To estimate the within-position principal angle, trials from each condition were randomly split in half. A subspace was estimated from each half, and the principal angle between the two halves was then computed. These procedures were repeated 30 times, and the resulting principal angles were averaged. The above analyses were performed at each encoding time point for each participant, and the estimates from the first 500 ms were further averaged.

### Ratio of variance explained (RVE)

Complementing principal angle, RVE was calculated to quantify the variance explained when data were projected onto a subspace (Li & Curtis, 2023), which resembles the variance accounted for (VAF) ratio (Xie et al., 2022). For instance, let S_1_ denote the sensor data of the conditions of interest, T_1_ the subspace estimated from **S**_1_, and **T**_2_ the subspace estimated from another set of sensor data of the conditions of interest. The RVE of S_1_ projected onto T_2_ was calculated as Var(**S**_1_**T**_2_)/Var(**S**_1_**T**_1_). Ultimately, RVE was quantified as the average of the RVE of S_1_ projected onto T_2_ and the RVE of S_2_ projected onto **T**_1_. For within-position estimates, data from each condition were randomly split into two halves, and results from 30 repetitions were averaged. The above procedures were performed at each encoding time point, and the estimates of the first 500 ms were averaged.

### Representational similarity analysis (RSA)

At each encoding time point, Spearman’s rank correlation was conducted on preprocessed signals of sensors (GP or LP sensors), utilizing spatial patterns as features. For position-specific representations (Figure 5C and Figure S3), trial pairs from different runs were categorized into four groups to compute spatial pattern similarity between trial pairs at the same global, local, and serial positions versus those at different positions.

### Power law exponent (PLE)

In terms of temporal dynamics, neural activity spans a broad range of frequencies, with slower frequencies exhibiting greater power than faster ones, reflecting scale-free properties. This phenomenon can be quantified by PLE. We calculated PLE using a windowed Fast Fourier Transform (FFT) on preprocessed sensor data, following the procedure described in a previous study (Wolff et al., 2019). For each participant, time series from a given period (i.e., the video-watching stage or the fixation period preceding video presentation) were concatenated across trials. The FFT was computed using a 20-s window with 50% overlap. The averaged FFT power spectrum was normalized by the total power across the entire frequency range. A log-log transform was then applied to the averaged FFT, with frequency (0-100 Hz) on the x-axis and power on the y-axis. A linear fit was applied to the transformed spectrum, and the absolute slope of the fitted line was defined as the PLE.

### Autocorrelation window (ACW)

The ACW was defined as the full-width-at-half-maximum of the autocorrelation function of the MEG time course (Honey et al., 2012). According to the Wiener-Khinchin theorem, the ACW and PLE are expected to represent related aspects of temporal structure. For each participant, preprocessed sensor signals from the same period were concatenated and then segmented into 20-s blocks with 50% overlap. The autocorrelation function, R_i_(τ), of the power of the i-th sensor within each block was computed as follows:

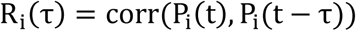

R_i_(τ) across all data blocks were then averaged, and the ACW of the i-th sensor was obtained using the following equation:

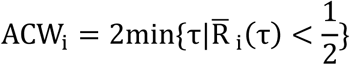

### Multidimensional distance (MDD)

We used MDD to quantify event-related representational changes between pre- and post-boundary periods, following a modified approach from Zheng et al. (2022). For each participant and each sensor group, the pre-boundary neural space was characterized using PCA (see Subspace analysis). **S**_pre_ was constructed from the averaged 400-ms pre-boundary data. The neural trajectories (i.e., time series of PC scores) for the three boundary conditions from 0 to 800 ms post-boundary were obtained by multiplying the trial-averaged data **S**_post_(t) at each time point t with the neural space **T**_pre_, which was estimated from **S**_pre_. This reflects the changes between post- and pre-boundary neural patterns. MDD was computed as the Euclidean distance between a given post-boundary time t and the boundary time point in the PC space (Figure 6B). The MDDs were averaged across the 0-800 ms post-boundary interval for further statistical comparisons (Figure 6C).

### Statistical analysis

#### Permutation test

To test the difference in the principal angle and the RVE, we randomly shuffled the condition labels of the observed data and computed the mean difference between the two conditions. This procedure was repeated 10,000 times to generate a null distribution. A p-value was then estimated by comparing the true mean difference to this null distribution. Note that between-position estimates of principal angle and RVE (e.g., global position 1 vs. 2) require comparisons using the corresponding within-position estimates (e.g., global position 1 vs. 1 and global position 2 vs. 2).

#### Cluster-based nonparametric permutation

To test decoding accuracy and position-specific representations across multiple time points, a nonparametric statistical test based on cluster-level permutation was conducted to control for multiple comparisons (Maris & Oostenveld, 2007). Specifically, for each time point, a t-test was first performed between the two conditions (e.g., same vs. different global positions), and neighboring elements with significant t-values were grouped into clusters. The summed t-score of all elements within each cluster was then computed. To assess significance, a null distribution of cluster-level statistics was generated by randomly permuting condition labels. For each permutation, the cluster with the maximal summed t-score was extracted, and this process was repeated 10,000 times to construct the null distribution. The true clusters were then compared against this distribution to compute their p-values.

