## supplementary material for "Hierarchical coding of local and global positions in episodic memory for naturalistic sequences"

This file includes:

Table S1. Statistical results for simple effect tests in a 2 (experimental groups) × 9 (serial positions) mixed design ANOVA

Figure S1. Decoding results for each position

Figure S2. Ratio of variance explained (RVE)

Figure S3. Serial position representation

Figure S4. Power percentage and autocorrelation window (ACW)

**Table S1. Statistical results for simple effect tests in a 2 (experimental groups) × 9 (serial positions) mixed design ANOVA**

| Position | $F_{(1,57)}$ | Partial $\eta^2$ |
|---|---|---|
| 1 | 5.74 | 0.0922 |
| 2 | 15.72 | 0.2158 |
| 3 | 12.54 | 0.1810 |
| 4 | 37.48 | 0.3966 |
| 5 | 24.04 | 0.2976 |
| 6 | 40.18 | 0.4132 |
| 7 | 69.82 | 0.5503 |
| 8 | 7.85 | 0.1199 |
| 9 | 0.44 | 0.0068 |

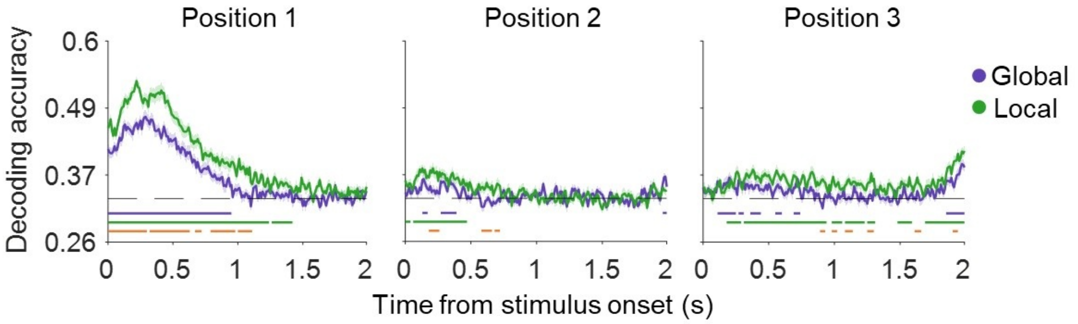


**Figure S1. Decoding results for each position**

Temporal dynamics of decoding accuracy for each global position (purple) and local position (green), with respective colored lines below the dashed lines (chance level) denoting significant temporal clusters of decoding accuracy. Orange lines indicate higher decoding accuracy for local than global positions. Shaded error bars represent the standard error of the mean across subjects.

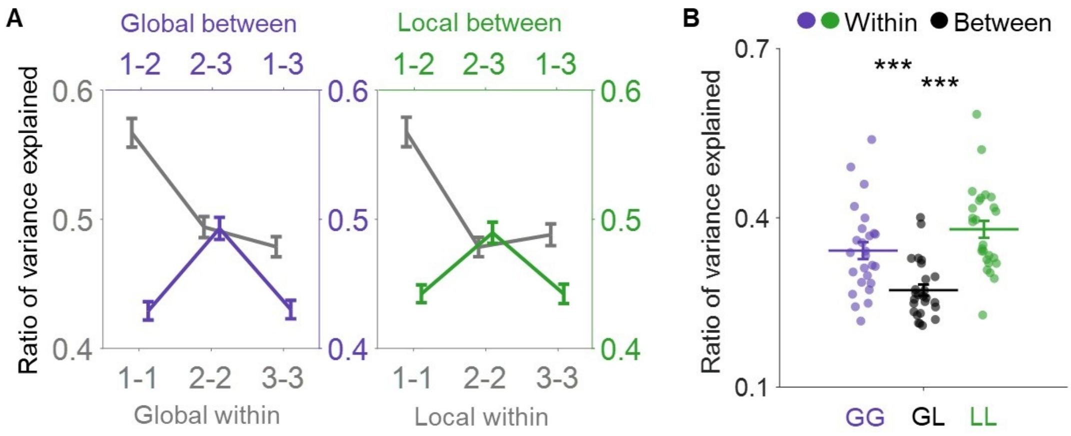


**Figure S2. Ratio of variance explained (RVE)**

(A) RVE for between-global (purple), within-global (gray), between-local (green), and within-local (gray) position subspaces. Error bars indicate the standard error of the mean across subjects.

(B) RVE for within-global (purple, GG), within-local (green, LL), and between-global-and-local (black, GL) position subspaces. Error bars indicate the standard error of the mean across subjects. Permutation test, ***$p < 0.001$.

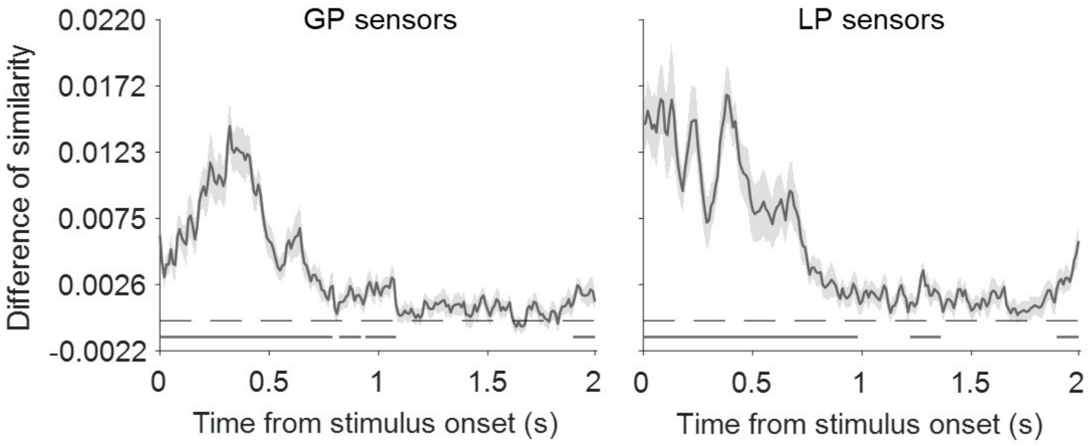


**Figure S3. Serial position representation**

Temporal clusters showing significant serial position representation are indicated by gray lines. Shaded error bars represent the standard error of the mean across participants.

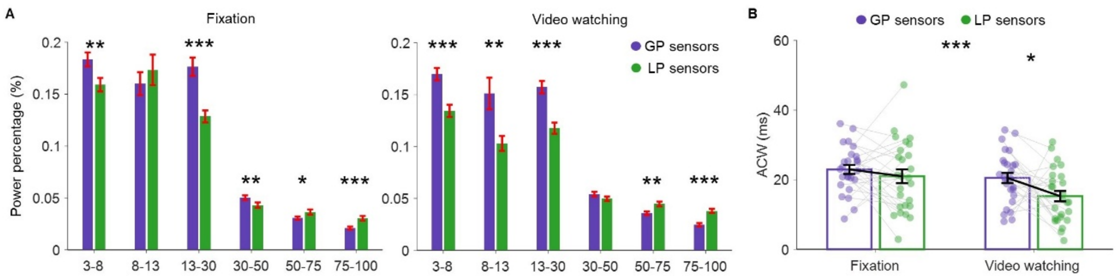


**Figure S4. Power percentage and autocorrelation window (ACW)**

(A) Power percentage across frequency bins for GP (purple) and LP (green) sensors. Error bars indicate the standard error of the mean across participants. *p<0.05, **p<0.01, ***p < 0.001.

(B) ACW for GP (purple) and LP (green) sensors. Error bars indicate the standard error of the mean across participants. *p < 0.05, ***p < 0.001.
